# Malassezia responds to environmental pH signals through the conserved Rim/Pal pathway

**DOI:** 10.1101/2024.07.11.603086

**Authors:** Kaila M. Pianalto, Calla L. Telzrow, Hannah Brown Harding, Jacob T. Brooks, Joshua A. Granek, Eduardo Gushiken-Ibañez, Salomé LeibundGut-Landmann, Joseph Heitman, Giuseppe Ianiri, J. Andrew Alspaugh

**Affiliations:** Departments of Medicine, Duke University School of Medicine, Durham, NC, USA; Molecular Genetics and Microbiology, Duke University School of Medicine, Durham, NC, USA; Biostatistics and Bioinformatics, Duke University School of Medicine, Durham, NC, USA; Pharmacology and Cancer Biology, Duke University School of Medicine, Durham, NC, USA; Cell Biology, Duke University School of Medicine, Durham, NC, USA; Department of Physics and Astronomy, University of North Carolina, Chapel Hill, NC, USA; Department of Agricultural, Environmental and Food Sciences, Università degli Studi del Molise, Italy; Section of Immunology at Vetsuisse Faculty, University of Zurich, Switzerland; Institute of Experimental Immunology, University of Zurich, Switzerland

## Abstract

During mammalian colonization and infection, microorganisms must be able to rapidly sense and adapt to changing environmental conditions including alterations in extracellular pH. The fungus-specific Rim/Pal signaling pathway is one process that supports microbial adaptation to alkaline pH. This cascading series of interacting proteins terminates in the proteolytic activation of the highly conserved Rim101/PacC protein, a transcription factor that mediates microbial responses that favor survival in neutral/alkaline pH growth conditions, including many mammalian tissues. We identified the putative Rim pathway proteins Rim101 and Rra1 in the human skin colonizing fungus *Malassezia sympodialis*. Gene deletion by transconjugation and homologous recombination revealed that Rim101 and Rra1 are required for *M. sympodialis* growth at higher pH. Additionally, comparative transcriptional analysis of the mutant strains compared to wild-type suggested mechanisms for fungal adaptation to alkaline conditions. These pH-sensing signaling proteins are required for optimal growth in a murine model of atopic dermatitis, a pathological condition associated with increased skin pH. Together these data elucidate both conserved and phylum-specific features of microbial adaptation to extracellular stresses.

**Importance:** The ability to adapt to host pH has been previously associated with microbial virulence in several pathogenic fungal species. Here we demonstrate that a fungal-specific alkaline response pathway is conserved in the human skin commensal fungus *Malassezia sympodialis* (*Ms*). This pathway is characterized by the pH-dependent activation of the Rim101/PacC transcription factor that controls cell surface adaptations to changing environmental conditions. By disrupting genes encoding two predicted components of this pathway, we demonstrated that the Rim/Pal pathway is conserved in this fungal species as a facilitator of alkaline pH growth. Moreover, targeted gene mutation and comparative transcriptional analysis supports the role of the *Ms* Rra1 protein as a cell surface pH sensor conserved within the basidiomycete fungi, a group including plant and human pathogens. Using an animal model of atopic dermatitis, we demonstrate the importance of *Ms* Rim/Pal signaling in this common inflammatory condition characterized by increased skin pH.

## Introduction

Changes in temperature, nutrient availability, or environmental pH create stressful conditions for microorganisms requiring continuous cellular adaptation for survival. In the case of pathogenic organisms, the shift from the ambient environment to the human host results in changes in many of these conditions, in addition to exposure to the host immune system. Similarly, commensal microorganisms must adapt to the unique environmental stresses presented by their specific niches within the host.

Extracellular pH can vary widely as microbes move from the environment to the human host. Even within the human body, pH can vary from the acidic pH of the stomach, to the more neutral pH of blood, to the basic pH of bile. Human skin tends to be more acidic than blood or other body sites, with pH levels that can vary from pH 4 to pH 6 on healthy adult skin [1]. This is an ideal pH for optimal growth of many fungi including *Malassezia sympodialis* (*Ms*) [2], a skin commensal microbe and opportunistic pathogen. However, in inflammatory skin conditions such as atopic dermatitis, the skin pH increases, representing an environmental trigger for this yeast-like species [2]. Additionally, other *Malassezia* species are adapted for colonization of the human gut. Recent reports reported that *Malassezia* can migrate from the more acidic upper gastrointestinal tract to more alkaline micro-niches in the pancreas, demonstrating how certain fungal species can adapt to fluctuations of pH within a mammalian host [3].

The fungus-specific Rim/Pal pathway includes a conserved cascading series of interacting proteins that sense and respond to changes in pH [4–6]. First described in model ascomycete fungi such as *Saccharomyces cerevisiae* and *Aspergillus nidulans*, this signaling pathway initiates a response to pH changes at the cell surface through the Rim21/PalH pH sensor [7–9]. This signal is then propagated through a conserved signaling cascade, with eventual cleavage and activation of the Rim101/PacC zinc-finger transcription factor [10–12]. Once activated, this transcription factor translocates to the nucleus where it regulates the expression of many genes, resulting in an adaptive cellular response to alkaline pH stress [6, 10, 13, 14]. By convention, this pathway has been referred to as the Pal pathway in filamentous fungi, and as the Rim pathway in fungi that grow predominantly as yeasts, such as *Saccharomyces cerevisiae*, *C. albicans*, and *C. neoformans*. Studies of this pathway in pathogenic fungi have revealed that many virulence factors require Rim/Pal pathway activation in order to be expressed. For example, in *Candida albicans*, the Rim pathway is required for the yeast-hyphal transition that is necessary for tissue invasion [15, 16]. Additionally, in the basidiomycete fungus *Cryptococcus neoformans*, activation of the Rim pathway is required for full expression of many virulence-associated phenotypes, including induction of the polysaccharide capsule and the formation of titan cells [10, 14, 17, 18].

Many Rim/Pal pathway proteins are conserved between the ascomycete fungi and the basidiomycete fungi *Cryptococcus neoformans* and *Ustilago maydis*, such as the components of the proteolysis complex (Rim20/PalA, Rim23/PalC, and the Rim13/PalB protease) and the Rim9/PalI chaperone [18–20]. Homologs of more upstream components comprising the pH-sensing complex, including the Rim21/PalH pH sensor and the Rim8/PalF arrestin, are notably absent in the genomes of basidiomycetes [18, 19]. The *C. neoformans* Rra1 protein was recently identified as a cell surface-associated protein that acts upstream of the Rim proteolysis complex to activate this pathway. While lacking sequence homology with the ascomycete Rim21 protein, *Cn* Rra1 shares functional and structural similarities with this established pH sensor [18] . Importantly, while Rim21 homologs are absent from the genomes of many basidiomycete fungi, Rra1 homologs are readily apparent in sequenced basidiomycete genomes [18]. We also identified a novel Rra1 interactor, Nucleosome Assembly Protein 1 (Nap1), which is required for activation of the Rim pathway in *C. neoformans*.

However, Nap1 homologs do not appear to be similarly involved in the Rim pathway of *S. cerevisiae* [21]. We therefore explored the degree of functional conservation of potential Rim pathway proteins in other basidiomycetes, such as *M. sympodialis*.

In this study, we established that the *M. sympodialis* Rim101 transcription factor and the putative Rra1 pH sensor are each required for survival at alkaline pH. In this way, we demonstrated that Rra1 is likely a conserved, basidiomycete-specific Rim pathway component. We also examined the transcriptional output of the *M. sympodialis* Rim pathway at alkaline pH, giving insight into the cellular processes involved in *M. sympodialis* survival in this stressful condition. Finally, we validated the relevance of the *Ms* Rim pathway in the interaction of this commensal fungus with the host by examining the innate immune response to fungal challenge *in vitro* and by evaluating the fitness of the fungus in the atopic skin environment *in vivo*.

## Materials and Methods

### Strains, media, and growth conditions

Strains used in this study are listed in **Supplemental Table S1**. Strains were routinely grown on modified Dixon’s (mDixon) medium (3.6% malt extract (Bacto), 1% mycological peptone (Oxoid), 1% ox bile (HiMedia), 1% v/v Tween 60 (Sigma), 0.4% glycerol; 2% Bacto agar added for plates) [22, 23]. For mDixon medium that was adjusted to specific pH, 150 mM HEPES was added to the medium as a buffering agent, and the pH was adjusted by addition of either concentrated HCl or NaOH. For *in vivo* experiments, strains were grown in a differently modified Dixon medium containing 3.6% malt extract (Sigma), 2% ox bile (Sigma), 0.6% bacterial peptone (Oxoid), 1% Tween 40 (Sigma), 0.2% glycoreol (Sigma) and 0.2% oleic acid (Sigma) [24]. Strains were cultured at 30°C unless otherwise indicated. *Agrobacterium tumefaciens* cultures were maintained on YT agar or FB broth (YT agar: 0.8% Bacto tryptone, 0.5% Yeast Extract, 0.5% NaCl, 1.5% glucose; FB broth: 2.5% Bacto tryptone, 0.75% yeast extract, 0.1% glucose, 0.6% NaCl, 50 mM Tris-HCl pH 7.6) at 30°C.

### *Agrobacterium tumefaciens*-mediated transformation

The *M. sympodialis RIM101* and *RRA1* genes were identified in the *M. sympodialis* genome assembly via BLASTp analysis [25]. To generate *M. sympodialis mutants*, targeted deletion constructs consisting of 1.5 kb of genomic sequence upstream and downstream the target genes and the *NAT* resistance marker were cloned into the T-DNA regions of plasmid pGI3 [26] using in vivo recombination in *Saccharomyces cerevisiae* as previously described [23]. Primers used to create the deletion constructs can be found in **Supplemental Table S2**. The resulting plasmid was transformed into *A. tumefaciens* EHA105 via electroporation and subsequent selection on YT agar supplemented with 50 µg/mL kanamycin.

*Agrobacterium*-mediated transformation was performed as described previously with some modifications [23, 27]. Briefly, *M. sympodialis* ATCC 42132 was incubated for 2 days at 30°C in mDixon liquid medium, and *A. tumefaciens* strains containing pKP38 (*RIM101* deletion) or pKP39 (*RRA1* deletion) were incubated overnight in FB + kanamycin at 30°C. *A. tumefaciens* cultures were diluted to an OD_600_ of 1 in Induction Medium (IM) (335) and incubated a further 4 hours at 30°C. *M. sympodialis* cells and induced *A. tumefaciens* cells were mixed at a 5:1 *Malassezia* : *Agrobacterium* ratio and pelleted at 5000 rpm, 10 min. The resulting cell pellet was spotted onto nylon membranes on modified IM (mIM) plates (IM medium supplemented with 0.4% ox bile, 0.4% v/v Tween 60 (Sigma), 0.1% v/v Tween 20) and incubated at RT for 7 days. After 7 days, cells were scraped from the membranes into sterile H_2_O and pelleted at 3000 rpm for 5 minutes (to preferentially pellet the *Malassezia* cells over the *Agrobacterium* cells). The resulting pellets were resuspended in sterile H_2_O and spread onto mDixon medium with cefotaxime + either nourseothricin or neomycin (300 µg/mL cefotaxime; 100 µg/mL nourseothricin; 200 µg/mL neomycin G418).

Resulting colonies were screened by PCR to confirm homologous recombination of the gene deletion constructs into the endogenous locus using primers listed in **Supplemental Table S3** [21]. Independent mutant strains for each gene were made by separate, completely independent transformations. All tested phenotypes were concordant between independent mutant strains; therefore, results from single or both mutants for each gene are demonstrated in the Results.

### Quantitative reverse-transcriptase PCR

As a pilot experiment prior to RNA sequencing, *M. sympodialis RIM101* expression was measured via RT-qPCR, per previous methods [21]. *M. sympodialis* was incubated overnight in mDixon at pH 4. Cells were pelleted by centrifugation and washed twice in sterile H_2_O, then resuspended in mDixon medium, mDixon buffered to described pH levels, RPMI + 10% FBS, or DMEM + 10% FBS. *M. sympodialis* was incubated in these media at 30°C for 90 minutes. At this time, *M. sympodialis* samples were pelleted by centrifugation for 5 minutes at 4000 rpm, 4°C. Cells were washed 2X with cold sterile H_2_O, then flash frozen on dry ice. RNA was purified using Trizol phenol-chloroform extraction (Invitrogen). cDNA was prepared using the AffinityScript cDNA synthesis kit (Agilent). qRT-PCR was performed using PowerUp SYBR Green (ThermoFisher). Relative *MsRIM101* gene expression levels compared to pH 4 conditions were calculated using the ΔΔC_T_ method with the *MsTUB1* tubulin gene as control [21]. RT-PCR primers for the *MsRIM101* and *MsTUB1* genes can be found in **Supplemental Table S4**.

### RNA Sequencing

To prepare samples for RNA sequencing, *M. sympodialis* WT, *rim101*Δ, and *rra1*Δ strains were grown in biological triplicate for 18 h in mDixon liquid medium at 30°C. Cultures were pelleted at 4000 rpm for 10 min, washed 1X with sterile H_2_O, and resuspended at an OD_6oo_ of 2.5 in 20 mL of mDixon pH 4 (Rim pathway-inactivating condition), mDixon pH 7.5 (Rim pathway-activating condition), or DMEM supplemented with 10% FBS (“host-like” condition). Cultures were incubated at 30°C for 90 minutes with shaking. Cultures were harvested by centrifugation at 4000 rpm for 10 min at 4°C, then washed once with cold sterile H_2_O. Cell pellets were flash frozen in a dry ice-ethanol bath and lyophilized overnight. RNA was extracted from the lyophilized pellets via a Trizol-chloroform extraction (Invitrogen). The resulting RNA was treated with DNase I (New England Biolabs) for 30 min at 37°C as described, then re-purified using Trizol and chloroform.

RNA Sequencing was performed in collaboration with the Duke University Center for Genomic and Computational Biology Sequencing and Genomic Technology Shared Resource. An mRNA library was prepared using a Kapa Stranded mRNA-Seq library prep kit. Stranded mRNA-Seq was performed on an Illumina HiSeq 4000 with 50-bp single-end reads. Reads were mapped to the *M. sympodialis* ATCC 42132 reference genome (obtained from Ensembl, accessed March 2021) using STAR alignment software [28]. Differential expression analyses were performed in R using a RNA-Seq Bioconductor workflow [29], followed by the DESeq2 package [30]. Genes were considered statistically differentially expressed if they had an adjusted P value [false-discovery rate (FDR)] of <0.05. Functional prediction for differentially regulated genes (+/- 1 log2 fold change) was performed using Fungi DB (www.fungidb.org [Release 63]) [31] entering the MSYG gene identifier for each transcript of interest.

The test of correlation between fold change values for *rra1*Δ and *rim101*Δ mutants was calculated using Kendall’s τ test, with the alternative hypothesis that fold change values were positively correlated. We initially only considered genes with significantly differential expression (adjusted p-value <= 0.05) in both mutants relative to WT. Results for among genes with significant differences in expression, using Kendall’s τ, of gene expression in between the growth at pH 7.5, in DMEM, and at pH 4 were, respectively, τ= 0.8206107, p-value < 2.2x10-16; τ= 0.71747, p-value < 2.2x10-16; τ= 1, p-value = 0.1666667). Because there are only 3 genes that have significantly different expression in at pH 4 (in both *rra1*Δ and *rim101*Δ mutants compared to WT), we repeated this calculation for all genes and again found significant results (with less dramatic τ values) for growth at pH 7.5 and in DMEM, but, again, not at pH 4 ((respectively τ= 0.6022764, p-value < 2.2x10-16; τ= 0.5515425, p-value < 2.2x10-16; τ= - 0.0465178, p-value = 0.999999). This analysis used the R packages: readxl, dplyr, ggplot2, and patchwork.

### Fungal survival in macrophages

The ability of the fungal strains to survive in the presence of macrophages was assessed by co-culture as previously described, with some alterations to accommodate *M. sympodialis* growth requirements [32] [33]. Approximately 10^5^ J774A.1 murine macrophages suspended in DMEM (Thermo Fisher Scientific) were added to individual wells of a 96-well plate and incubated overnight at 37°C with 5% CO_2_. Following adherence to the 96-well plate, J774A.1 murine macrophages were activated with 10 nM phorbol myristate acetate (PMA) in RPMI 1640 medium (Corning) supplemented with 20% FBS for 1 hour at 37°C with 5% CO_2_. Following macrophage activation, *M. sympodialis* strains (WT, *rim101*Δ mutant strains [KPY34 and KPY36], and *rra1*Δ mutant strains [KPY38 and KPY39]), which had been incubated for 48 hours in mDixon medium, were washed three times in sterile water, normalized to an OD of 0.3 in RPMI 1640 medium supplemented with 20% FBS, and added to the activated J774A.1 murine macrophages (5 x 10^6^ fungal cells per well (10:1 fungal cells:macrophages)). Co-cultures of J774A.1 murine macrophages and fungal cells were incubated for 4 hours at 37°C with 5% CO_2_. Phagocytosed fungal cells were collected by washing individual wells of the 96-well plate vigorously with sterile water. Collected fungal cells were plated onto mDixon agar to assess the number of viable *M. sympodialis* cells by quantitative culture. The results are reported as the average percentage (± SEM) of recovered CFU, normalized to the WT strain, generated from at least 3 biological replicates. Statistical significance was determined using one-way analysis of variance (ANOVA) and the Tukey-Kramer test (GraphPad Software, San Diego, CA).

Supplementation with 20% FBS provided exogenous lipids to support *M. sympodialis* lipid auxotrophy in this assay. RPMI 1640 medium was utilized specifically in this experiment to limit the impact of alkaline pH on the survivability of the *rim101*Δ and *rra1*Δ mutant strains. We found that a 4-hour incubation in RPMI 1640 medium supplemented with 20% FBS without J774A.1 murine macrophages did not impact the survivability of the *rim101*Δ and the *rra1*Δ mutant strains compared to the WT strain. Attempts at longer incubations, such as 24 hours, failed to recover viable fungi from co-culture. As a result, we used 4-hour co-cultures to directly assess the ability of the tested fungal strains to interact with and survive in the presence of macrophages.

### Macrophage activation assays

Bone marrow cells were isolated from C57BL/6 mice (Jackson Laboratories) as previously described [34, 35]. Briefly, femurs were isolated from CO_2_-euthanized mice, and each bone marrow space was flushed with cold PBS. Red blood cells were lysed in 1x RBC lysis buffer, and the remaining bone marrow cells were resuspended in 1x Dulbecco’s modified Eagle’s medium (DMEM) with 1 U/ml penicillin/streptomycin (PenStrep). Adherent cells were differentiated in BMM medium (1x DMEM, 10% fetal bovine serum [FBS; non-heat inactivated], 1 U/ml penicillin/streptomycin) with 3 ng/ml recombinant mouse GM-CSF (rGM-CSF; R&D Systems or BioLegend) at a concentration of 2.5 x 10^5^ cells/ml in 150 x 15 mm petri plates at 37°C with 5% CO_2_. The media was refreshed after 3 days and the cells were harvested on day 7 as previously described, likely resulting in a mixture of bone-marrow-derived macrophages (BMMs) and dendritic cells (DCs) [35].

These cells were counted (by hemocytometer, with Trypan blue to differentiate between live and dead cells), plated in BMM medium in 96-well plates at a concentration of 5 x 10^4^ cells per well, and incubated at 37°C with 5% CO_2_ overnight prior to fungal co-culture experiments.

BMM co-cultures with wildtype *M. sympodialis*, *C*. *neoformans,* and *C. albicans* as well as *M. sympodialis rim101*Δ and *rra1*Δ mutant strains were performed as described previously [34, 35]. *M. sympodialis* strains (WT (ATCC 42132), *rim101*Δ (KPY34), and *rra1*Δ (KPY36)) were incubated for 3 days in DMEM media supplemented with 10% fetal bovine serum [FBS; non-heat inactivated] and 1 U/ml penicillin/streptomycin) at 30°C. The *C. neoformans* H99 strain was incubated for 2 days in DMEM media supplemented with 10% fetal bovine serum [FBS; non-heat inactivated] and 1 U/ml penicillin/streptomycin) at 30°C. Prior to co-culturing with BMMs, these cultures were transferred to 37°C for 16 hours. WT *C. albicans* (SC5314) cells were incubated for 1 day in DMEM without serum at 30°C prior to co-culture to avoid premature filamentation. Following these incubations, fungal cells were washed twice with PBS, counted, and added to wells of a 96-well plate containing BMM (5 x10^5^ BMM/well) at a concentration of 5 x 10^6^ fungal cells per well (10:1 fungal cells:BMMs). Co-cultures were incubated for the indicated amount of time (either 3 or 6 hours) at 37°C with 5% CO_2_. Supernatants were collected and stored at -80°C overnight. Secreted cytokines (TNF) were quantified in supernatants by enzyme-linked immunosorbent assay (ELISA MAX:Deluxe Set Mouse TNF); BioLegend). Data are represented as the average TNF levels (pg/ml) for 4-5 biological replicates per group [34, 35].

### Scanning electron microscopy (SEM)

SEM was used to visualize the interactions between *M. sympodialis* strains and J774A.1 murine macrophages. Co-cultures were performed as described above, with some alterations. Individual ethanol-sterilized polydopamine coated coverslips (18 mm diameter) were placed into the wells of a 12-well plate. Approximately 5 x 10^6^ J774A.1 murine macrophages suspended in RPMI 1640 medium supplemented with 20% FBS and 10 nM PMA were added to each well and incubated for 1 hour at 37°C with 5% CO_2_. Following adherence and activation, 48-hour incubated *M. sympodialis* cultures (WT, *rim101*Δ mutant [KPY34], and *rra1*Δ mutant [KPY38]) were washed three times in sterile water, normalized to an OD of 0.3 in RPMI 1640 medium supplemented with 20% FBS, and added to wells (5 x 10^6^ fungal cells per well (1:1 fungal cells:macrophages). Co-cultures of J774A.1 murine macrophages and fungal cells were incubated for 1 hour at 37°C with 5% CO_2_ to capture potential interactions (such as phagocytosis) between macrophages and fungal cells.

Co-cultures were fixed with 2.5% glutaraldehyde for 1 hour at room temperature and were subsequently washed 3 times with 1X PBS. Samples were dehydrated by immersing the coverslips in ethanol (30% for 5 minutes, 50% for 5 minutes, 70% for 5 minutes, 95% for 10 minutes, and 100% for 10 minutes performed twice). Samples were then critical point dried with a Tousimis 931 critical point dryer (Rockville, Maryland) and coated with gold-palladium using a Cressington 108 sputter-coater (Watford, United Kingdom). Coverslips containing the prepared samples were mounted and imaged on a Hitachi S-4700 scanning electron microscope (Tokyo, Japan).

### Murine skin colonization assay

Mouse experiments in this study were performed in strict accordance with the guidelines of the Swiss Animals Protection Law and under protocols approved by the Veterinary office of the Canton Zurich, Switzerland (license number 142/2021). All efforts were made to minimize suffering and ensure the highest ethical and humane standards according to the 3R principles [36]. WT C57Bl/6JRj mice were purchased from Janvier Elevage (France). All experiments were conducted at the Laboratory Animal Science Center of the University of Zurich under specific pathogen-free conditions. AD-like conditions were induced in the murine ear skin according to Moosbrugger-Martiz et al. [37] and Ruchti et al. [38]. Briefly, 1.125 nm of MC903 (calcipotriol hydrate, Sigma) diluted in pure ethanol was applied on the dorsal and ventral side of both ears for 5 consecutive days and again for 4 days after a resting period of 2 days. The pH in MC903-treated mouse skin is higher (pH 7-7.5) compared to that of control skin (pH 5.5) [39]. *M. sympodialis* strains (WT, *rim101*Δ, and *rra1*Δ) were grown in mDixon for 2 days, washed twice in PBS and resuspended in native olive oil. A suspension of 100 μl containing 1x10^7^ yeast cells was applied topically onto the dorsal side of both ears while mice were anaesthetized [40]. After infection, MC903 treatment was continued on day 2 p.i. (for the 4 days infection experiment) or daily from day 2 to 5 (for the 7 days infection experiment) on the ventral side of the ear only to avoid interference of the EtOH solvent with fungal viability [38]. Ear thickness was continuously monitored using the Oditest S0247 0-5 mm measurement device (Kroeplin). For determining the fungal loads in the skin, the ear tissue was transferred in water supplemented with 0.05% Nonidet P40 (AxonLab), homogenized with a TissueLyzer (Qiagen) for 6 minutes at 25 Hz, plated on mDixon agar, and incubated at 30°C for 3 to 4 days for colony counting.

### Data Accessibility

Processed RNA-seq seq data are available in the supplemental Tables; raw and processed data are available through NCBI Gene Expression Omnibus (GEO) accession number GSE254653 (https://www.ncbi.nlm.nih.gov/geo/query/acc.cgi?acc=GSE254653).

## Results

### Mutants in *M. sympodialis* Rim pathway signaling are hypersensitive to elevated pH and salt concentrations

In other fungal species, the Rim/Pal pathway is involved in sensing and responding to increases in extracellular pH [4, 5, 14, 18]. Accordingly, Rim pathway genes are required for survival at elevated pH as well as at elevated salt concentrations [15, 17, 41–43]. To determine whether the *M. sympodialis* (*Ms*) Rim pathway is involved in similar cellular responses, two putative Rim pathway genes in the recently annotated *Ms* genome were identified based on sequence homology; MSYG_3336 encodes the closest homolog of the Rim101 transcription factor, and MSYG_4280 encodes the closest homolog of the Rra1 putative pH sensor. To explore their roles in the fungal response to extracellular stresses, two independent loss-of-function mutant strains were generated for each corresponding gene (**Supplemental Table S1**). All tested phenotypes were concordant between the independent *Ms rra1*Δ mutants as well as between the independent *Ms rim101*Δ mutants. On media containing exogenous lipids, the wild-type *Ms* strain was able to grow well from a pH of 5 to a pH of 7.5; the *Ms rim101*Δ and *rra1*Δ mutant strains grew at rates similar to the WT at mildly acidic pH (pH 6) (**Figure 1A**). However, they began to display growth defects at neutral pH 7, and growth was completely inhibited at pH 7.5 (**Figure 1A**). Similar to Rim/Pal pathway mutants in other fungi, these *Ms* mutant strains displayed a growth impairment at high concentrations of NaCl (**Figure 1A**). These results suggest that, as observed in other fungal genera, the *Ms* Rim101 transcription factor is involved in sensing and responding to alkaline pH and high salt conditions. Moreover, these data also suggest that Rra1 homologs play a conserved role in basidiomycete Rim/Pal pathways to sense and respond to alkaline pH signals.

**Figure 1.**
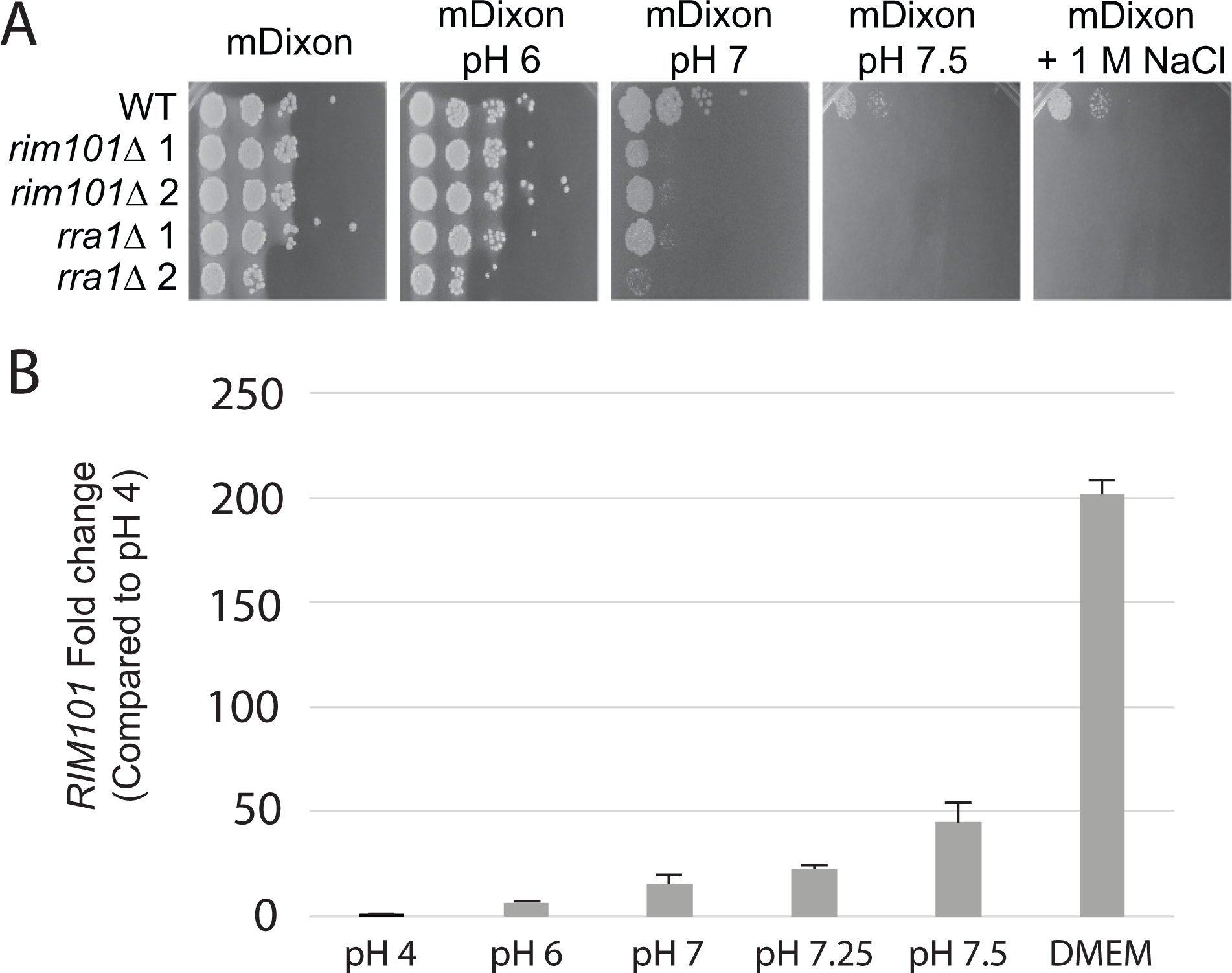
(A) ***M. sympodialis RIM101* and *RRA1* genes are required for optimal growth at alkaline pH and in high salt conditions**. Indicated strains were serially diluted and incubated in spot cultures on mDixon medium buffered to pH 6, pH 7, or pH 7.5, or mDixon medium supplemented with 1 M NaCl to examine growth phenotypes. Plates were imaged 6 days post-inoculation. (B) ***M. sympodialis RIM101* gene expression in response to increasing pH and tissue culture medium.** Wild-type *M. sympodialis* cells were incubated overnight in mDixon medium pH 4, then shifted to one of the following conditions for 90 minutes: mDixon buffered to pH 4, pH 6, pH 7, pH 7.25, or pH 7.5; or tissue culture medium (DMEM or RPMI + 10% FBS). Relative transcript abundance of the *RIM101* gene was assessed by quantitative real-time PCR using the ΔΔC_T_ method and the *TUB2* tubulin gene as control. Fold-change values for each condition were normalized to mDixon pH 4.

### Comparative transcriptional analysis defines Rim pathway-regulated genes in *M. sympodialis*

Activation of the conserved and fungal-specific Rim signaling pathway results in the proteolytic cleavage of the Rim101 transcription factor, which then translocates to the nucleus to regulate transcriptional responses to alkaline pH, including induction of the expression of *RIM101* itself [14]. To first determine pathway activating signals, we performed quantitative real-time PCR to assess conditions associated with transcriptional activation of the *RIM101* gene. We performed this transcriptional analysis across a gradient of pH values in mDixon medium, while also assessing *RIM101* transcript levels in tissue culture medium. *Ms RIM101* transcript levels increased in response to a more alkaline pH in a dose-dependent manner (**Figure 1B**), similar to *RIM101/PacC* transcriptional induction observed in other fungal species [44, 45]. Additionally, *RIM101* is even more strongly transcriptionally induced in tissue culture media (DMEM and RPMI media, pH 7.4) than mDixon medium pH 7.5, indicating that signals in tissue culture media in addition to pH are involved in activating the Rim pathway in this organism (**Figure 1B**).

We performed deep RNA sequencing comparing the transcriptomes of the *Ms* WT, *rim101*Δ, and *rra1*Δ strains incubated for 90 minutes in mDixon pH 4, mDixon pH 7.5, and DMEM tissue culture medium (pH 7.4). Gene expression is highly correlated between *rra1*Δ and *rim101*Δ mutants, compared to WT, when grown at pH 7.5 or in DMEM, but there is little correlation between *rra1*Δ and *rim101*Δ mutants when they are grown at pH 4 (Figure S1). This correlation is highly statistically significant at pH 7.5 or in DMEM (p-value < 2.2x10-16 for both), but there is no significant correlation for growth at pH 4 (p-value = 0.1666667).

Because there are only three genes that have significantly different expression at pH 4 in both *rra1*Δ and *rim101*Δ mutants compared to WT, we also calculated Kendall’s τ for all genes and again found significant results (also less dramatic) for growth at pH 7.5 and in DMEM, but not at pH 4 (respectively τ= 0.6022764, p-value < 2.2x10-16; τ= 0.5515425, p-value < 2.2x10-16; τ= -0.0465178, p-value = 0.999999)

We also analyzed the combined transcriptional data from each experimental sample using multidimensional scaling (MDS) analysis to visually compare variations in the transcriptomes of each dataset. As expected, biological replicates of the same strain incubated at the same conditions tended to cluster more closely to each other than to other samples (**Figure 2A**). This analysis indicates that overall patterns of transcriptional activity are very similar in mDixon medium at pH 4 among the WT, *rim101*Δ, and *rra1*Δ strains (**Figure 2A**), consistent with work in other fungal species demonstrating specific activation of the Rim/Pal pathway in response to alkaline pH and other cell stress signals [4]. In contrast, when these strains were incubated at elevated pH in mDixon medium buffered to pH 7.5 for 90 minutes, there are distinct patterns of transcriptional activity that distinguish the WT strain from the two mutant strains, which clustered together and clearly distinctly from the WT strain (**Figure 2A**). A similar observation was made for samples obtained after growth in DMEM, although there was a higher variation among the *rim101*Δ and *rra1*Δ mutants. The relative transcriptional variation of individual genes between the WT and the *rim101*Δ / *rra1*Δ strains at each pH is demonstrated by Volcano plots (**Figure 2B**).

**Figure 2.**
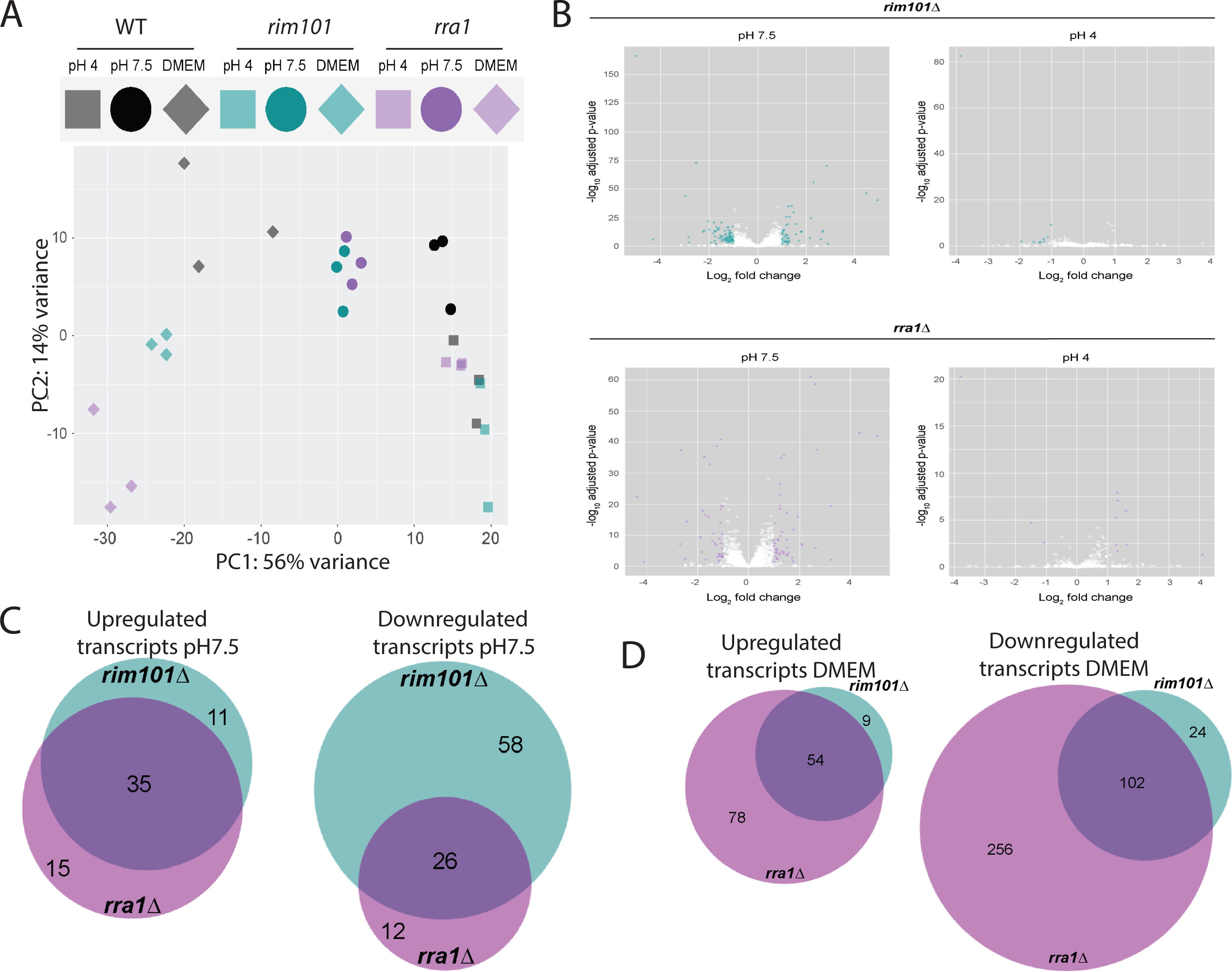
Comparative transcriptional analysis. The *M. sympodialis* WT, *rim101*Δ, and *rra1*Δ strains were incubated in each of the following conditions for 90 minutes prior to total RNA extraction for RNAseq analysis: mDixon medium pH 4, mDixon medium pH 7.5, DMEM. (**A**) Comparison of global transcriptome patterns by Multidimensional Scaling (MDS) analysis (3 biological replicates for each indicated strain in each condition). (**B**) Volcano plots illustrating number of genes with statistically significant alterations in transcript abundance at pH 7.5 and pH4. Genes with alterations in transcript abundance (+/- 1 log_2_ fold change) are indicated in green (*rim101*Δ versus WT) or magenta (*rra1*Δ versus WT). (**C**) Venn diagram indicating the number of genes with statistically significant differences in transcript abundance (+/- 1 log_2_ fold change) for indicated strains compared to WT at pH 7.5 (green = *rim101*Δ, magenta = *rra1*Δ). (**D**) Venn diagram indicating the number of genes with statistically significant differences in transcript abundance (+/- 1 log_2_ fold change) for indicated strains compared to WT in DMEM (green = *rim101*Δ, magenta = *rra1*Δ).

At pH 4, there are very few genes differentially regulated between the WT and *rim101*Δ or *rra1*Δ mutant strains (**Figure 2B; Tables S5, S6**). However, at pH 7.5, 84 *Ms* genes displayed statistically significant decreased transcript abundance in the *rim101*Δ mutant strain compared to WT (< -1 log_2_ fold change), suggesting their transcriptional dependence on the putative Rim101 transcription factor (**Figures 2B, 2C; Tables S7, S8**). Of these genes, 26 (31%) demonstrated similar decreases in transcript abundance in the *rra1*Δ mutant (**Figure 2C; Tables S7, S9, S8, S10, S11**). Additionally, 46 genes displayed increased transcript abundance (> +1 log_2_ fold change) in the *rim101*Δ strain compared to WT, and 35 (76%) of these also had increased transcript abundance in the *rra1*Δ strain (**Figures 2B, 2C; Tables S7, S12, S13, S11**).

Similar overlapping transcriptional patterns of the *Ms rim101*Δ and *rra1*Δ mutant strains are observed in DMEM tissue culture medium (pH 7.4): 126 *Ms* genes display decreased transcript abundance in the *rim101*Δ mutant strain compared to WT, and 102 (81%) of these genes also demonstrate decreased transcript abundance in the *rra1*Δ mutant; 63 genes have increased transcript abundance in the *rim101*Δ strain compared to WT, and 54 (86%) have similarly increased transcript abundance in the *rra1*Δ strain (**Figure 2D; Tables S14, S15, S16, S17, S18, S19, S20**).

Assigning likely function to specific genes in these datasets is limited by the incomplete annotation of the *M. sympodialis* genome (>20% of genes listed as encoding an “unspecified product”). However, we used FungiDB [31] to assist in assigning predicted function for the proteins encoded by genes with altered transcription in either the *rim101*Δ or *rra1*Δ mutant strains at pH 7.5 and in DMEM (**Tables S12, S10, S16, S17, S13, S10, S18, S19**). We also manually defined functional categories of predicted function for genes with similar patterns of altered transcription in both the *rim101*Δ and *rra1*Δ strains in these two incubation conditions. A subset of these genes is listed in **Tables 1 and 2**. This analysis suggested potential *Ms* Rim pathway regulation at alkaline pH for multiple genes encoding proteins involved in membrane transport (MFS proteins; transporters of ammonium, amino acids, nucleosides, and ions) and intracellular trafficking (including the ESCRT II protein Vps25). A larger gene set displayed altered transcript abundance in DMEM between the *rim101*Δ / *rra1*Δ strains and WT. These genes similarly included those predicted to encode proteins involved in membrane transport and intracellular trafficking. Additional functional categories for genes with differential transcript abundance in DMEM in these Rim pathway mutants included cell cycle regulation, lipid metabolism, and cell surface modification. These categories are similar to those for Rim pathway-regulated genes in related basidiomycetes in which cell surface (cell wall and membrane) and cell cycle modifications are important components of the adaptive cellular response to environments with elevated pH [45].

**Table 1.**
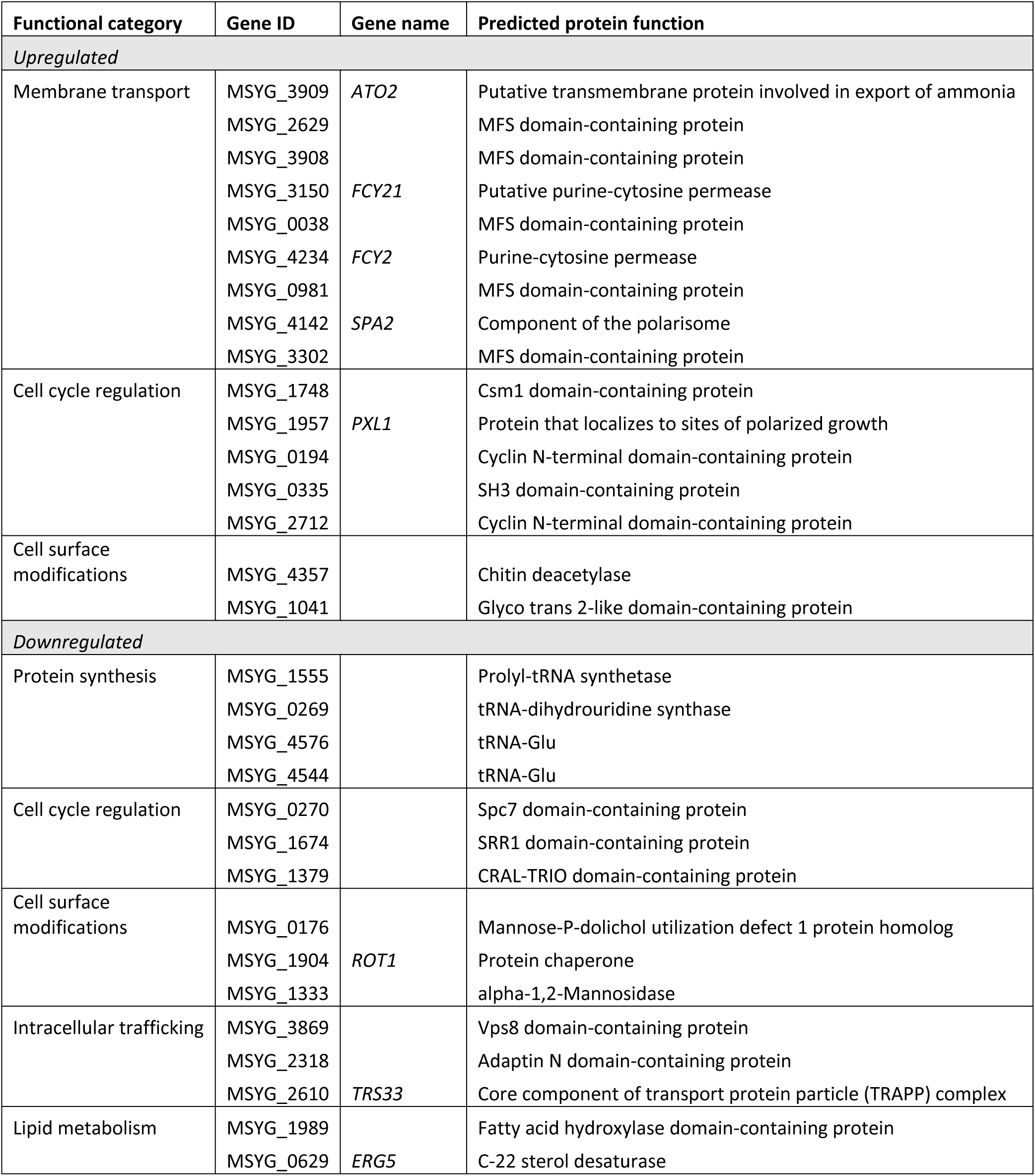
Functional categories enriched among the genes differentially expressed in DMEM in both the *rim101*Δ and *rra1*Δ mutant strains. Functional category assignment was performed manually based on predicted gene function using FungiDB annotations (2022).

**Table 2.**
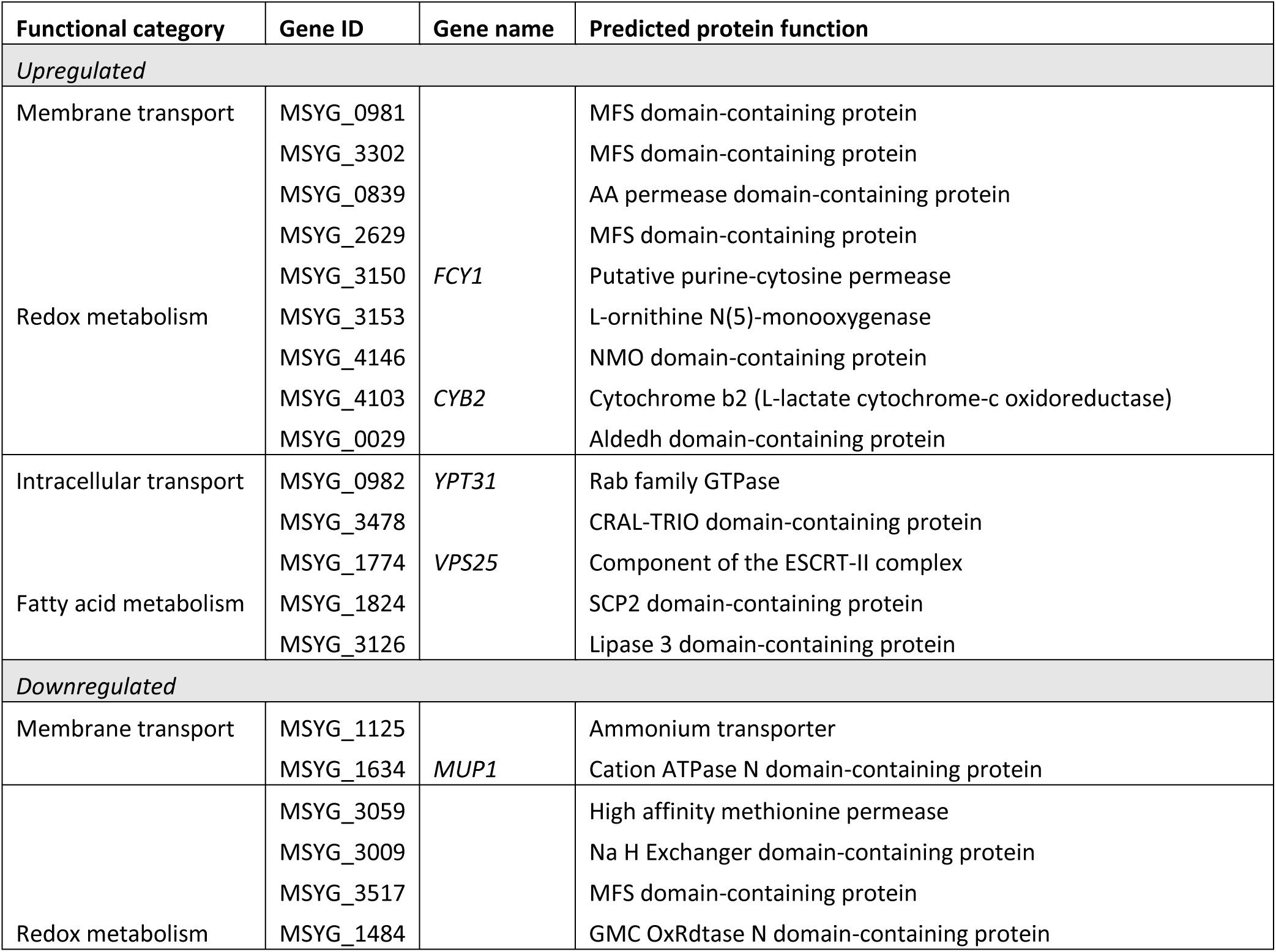
Functional categories enriched among the genes differentially expressed in the presence of alkaline pH (pH 7.5) in both the *rim101*Δ and *rra1*Δ mutant strains. Functional category assignment was performed manually based on predicted gene function using FungiDB annotations (2022).

The *Ms RRA1* gene displayed transcriptional dependence on the Rim101 transcription factor in DMEM (-0.95 log_2_ fold change). This suggests that potentially biologically relevant changes in the transcript abundance of Rim101-regulated genes might be missed by arbitrary fold-change cut-offs and subsequent dataset limitations.

### *M. sympodialis* interacts with macrophages *in vitro* in a Rim pathway-independent manner

Due to its prevalence on the skin, *Malassezia* species have been studied for interactions with epidermis-resident keratinocytes, dendritic cells, and macrophages [24] as well as with the murine macrophage-like cell line J774A.1 *in vitro* [33]. To assess the role of *M. sympodialis* Rim signaling in the interaction of fungal and innate immune cells, we performed a similar *in vitro* co-culture of the WT and Rim pathway mutants with J774A.1 cells. As early as one hour after co-culture, we observed macrophages clustering with fungal cells (**Figure 3A**). We used scanning electron microscopy (SEM) to further visualize the details of these physical interactions. We found that all tested strains (WT, *rim101*Δ [KPY34], and *rra1*Δ [KPY38]) were actively phagocytosed within one hour of co-culture (**Figure 3B**).

**Figure 3.**
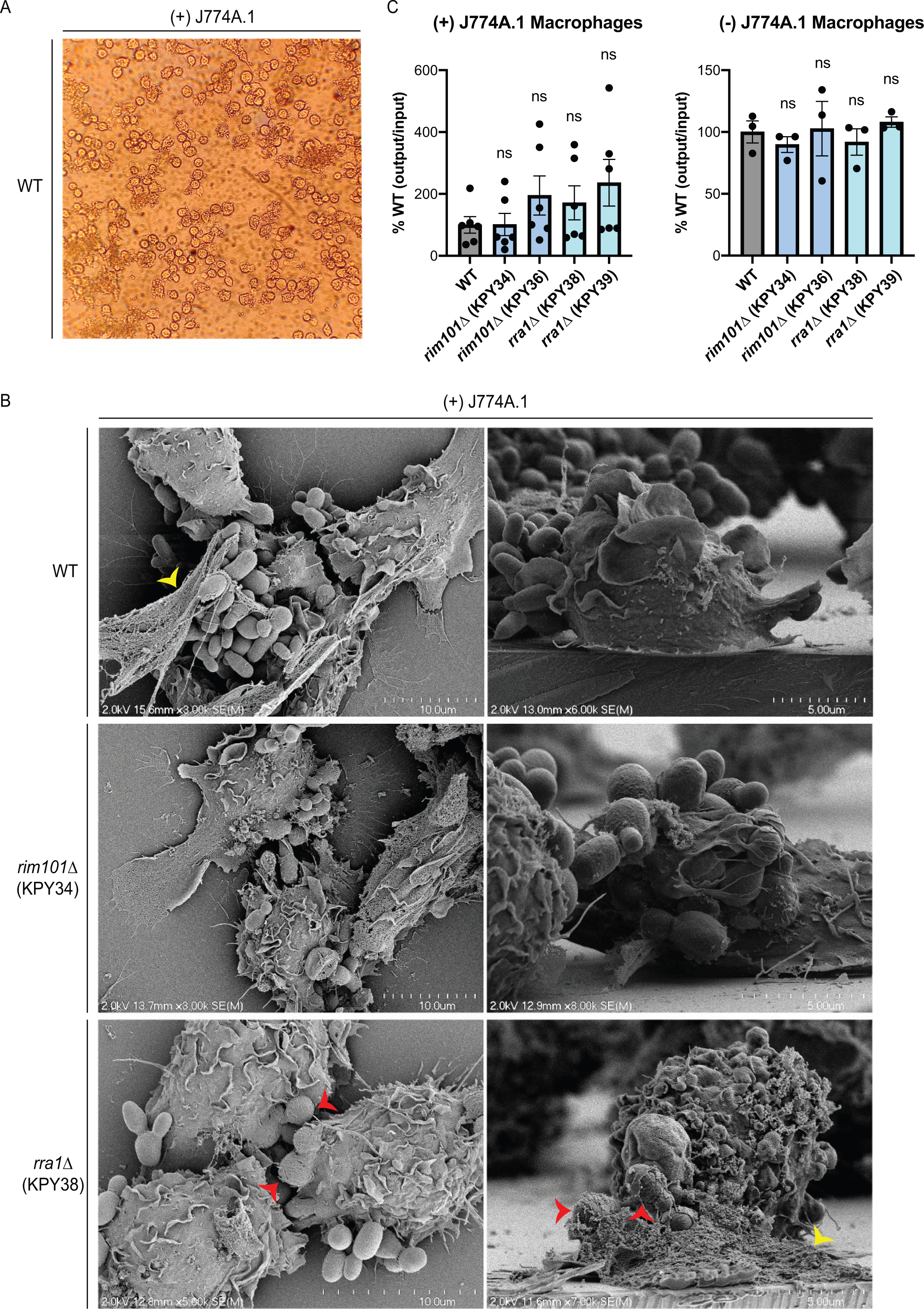
*M. sympodialis* interactions with macrophages. (**A**) Light microscopy image of the WT strain co-cultured with J774A.1 macrophages after one hour of co-culture. (**B**) The WT strain, *rim101*Δ mutant strains, and *rra1*Δ mutant strains were co-incubated with (left) and without (right) J774A.1 murine macrophages for 4 hours. Survival of the indicated fungal strains was assessed by quantitative culture, and the percentage of recovered (output) colony forming units (CFUs) compared to the original (input) CFUs was normalized to the WT strain [% WT (output/input)]. This experiment was performed with a minimum of three biological replicates (*n* = 3). Error bars represent the SEM compared to WT. Log transformation was used to normally distribute the data for statistical analysis (one-way ANOVA; ns, not significant). (**C**) Scanning electron microscopy (SEM) images of the WT strain, the *rim101*Δ mutant strain (KPY34), and the *rra1*Δ mutant strain (KPY38) after one hour of co-culture with J774A.1 macrophages. Cells were fixed with 2.5% glutaraldehyde, dehydrated, critical point dried, sputter-coated with gold-palladium, and imaged. Red arrowheads indicate fungal cells actively undergoing macrophage phagocytosis. Yellow arrowheads indicate potential macrophage extracellular traps (METs).

Typically, we observed groups of ∼2-3 macrophages assembling to engulf clumps of fungal cells. In some cases, the macrophage/fungal cell association contained extracellular material, possibly consistent with macrophage extracellular traps (METs) [46] (**Figure 3B**, yellow arrow heads).

To assess the ability of this macrophage-like cell line to rapidly kill the fungal cells, we also tested for survival differences among the three *M. sympodialis* strains in this co-culture system [33]. We chose a short co-incubation period to address pH-related growth effects in the *rim101*Δ and *rra1*Δ strains. After four hours of co-culture with J774A.1 cells, we observed no significant survival differences between the WT, *rim101*Δ, and *rra1*Δ mutants (**Figure 3C, left**). Despite the alkaline pH growth defect of the two mutants, there was no loss of viability of either mutant strain when incubated in tissue culture medium without macrophages during this short period of incubation (**Figure 3C, right**). Collectively, these observations indicate that macrophages recognize and actively phagocytose *Ms.* However, this association does not result in rapid fungal cell killing.

### *M. sympodialis* elicits a robust TNF response when co-cultured with macrophages independent of Rim pathway signaling

Prior investigations into the ability of *Malassezia* to activate macrophages have yielded conflicting results, with individual studies suggesting that *Malassezia* species may either trigger or inhibit macrophage activation. These discrepancies have been attributed to differences in the growth phase of the cells, the fungal species assayed, and the nature of the protective lipid layer surrounding *Malassezia* cells [47–49]. We quantified the production of tumor necrosis factor (TNF) following co-incubation of primary bone marrow-derived macrophages (BMMs) with the *M. sympodialis* wildtype, *rim101*Δ mutant, and *rra1*Δ mutant strains. We also measured TNF production in response to co-culture with wild-type *Candida albicans* and *Cryptococcus neoformans* strains, to serve as positive and negative controls, respectively. Following both a 3-hour and 6-hour co-culture (**Figure 4A, B**), we observed an expected profound induction of TNF by the macrophages co-cultured with *C. albicans* cells [50]. We also noted undetectable levels of TNF after co-culture with wildtype *C. neoformans* cells, consistent with prior studies documenting an effective immune evasion phenotype by wild-type *C. neoformans* cells [17]. Similar to the case with *C. albicans*, BMMs exposed to all three *M. sympodialis* strains produced high levels of TNF (**Figure 4A, B**). To determine whether this level of TNF by BMMs required viable *M. sympodialis* cells, we repeated the assay with heat-killed fungal cells. We observed similar levels of TNF production as with live cells in all strains tested (**Figure S2**), suggesting that the TNF production is most likely primarily due to a physical interaction between the macrophage and structural features of the fungal cell, as observed in other fungal species [34]. Exposure of the BMMs to the *Ms rra1*Δ mutant strain resulted in variably lower levels of TNF-stimulation compared to the WT and *rim101*Δ mutant, though not resulting in the complete suppression of macrophage activation as observed with the *C. neoformans* control (**Figure S2**) [17]. Together these data suggest that the cell changes associated with mutation of the *M. sympodialis RIM101* gene do not affect TNF production by co-cultured macrophages, and a *Ms rra1*D mutation has only a partial effect. Moreover, the degree of macrophage TNF production in response to *Ms* is similar to that observed in co-cultures with *C. albicans*. In contrast, *M. sympodialis* does not share the immune avoidance phenotype of *C. neoformans* in this assay.

**Figure 4.**
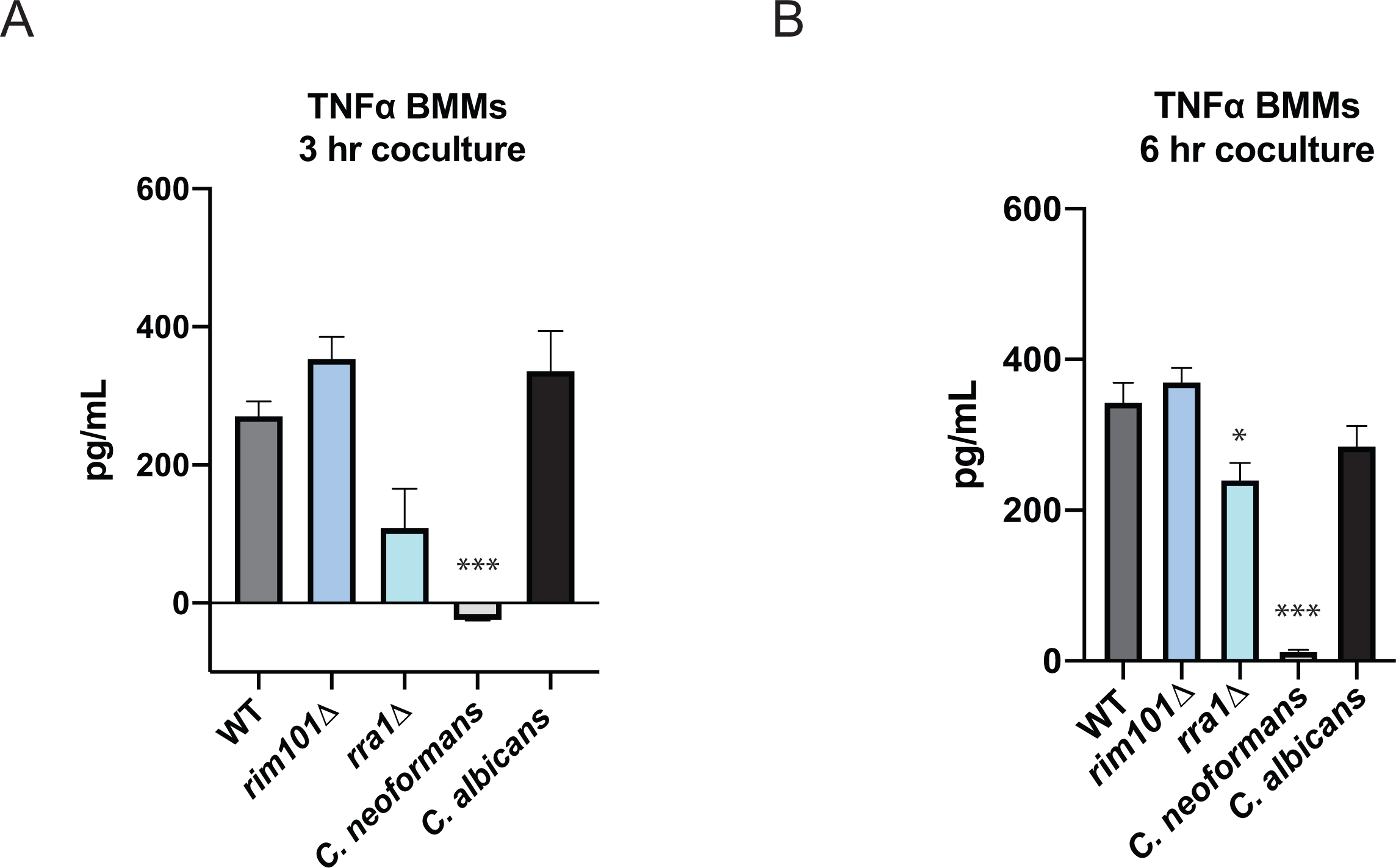
*M. sympodialis* elicits TNF response when co-cultured with macrophages. The *M. sympodialis* WT, *rim101*Δ, and *rra1*Δ strains were incubated for 3 days in DMEM + 10% FBS at 30°C preceding a final incubation at 37°C for 16 hours prior to co-culture. WT *C. neoformans* (strain H99) cells were incubated for 2 days in DMEM + 10% FBS at 30°C preceding a final incubation at 37°C for 16 hours prior to co-culture. WT *C. albicans* (SC5314) cells were incubated for 1 day in DMEM without serum at 30°C prior to co-culture. Bone marrow-derived macrophages (BMMs) were co-incubated with the indicated fungal strains for 3 hours (**A**) or 6 hours (**B**) at a multiplicity of infection (MOI) of 10:1, fungal cells:BMMs. TNF levels (pg/ml) were assayed from the co-culture supernatant by ELISA. Data represent means from 6 replicates per strain per condition. One-way ANOVA and Tukey’s multiple comparison test were used to compare means. ****, p < 0.0001; ***, p < 0.001; *, p = 0.0186. Statistical comparisons were made against WT *M. sympodialis*.

### The Rim/Pal pathway impacts *M. sympodialis* fitness in a murine model of atopic dermatitis

Finally, we assessed the impact of the Rim/Pal pathway on the interaction of *M. sympodialis* with the host skin *in vivo* under high pH conditions reminiscent of those in the skin of atopic dermatitis patients [39]. Atopy-like conditions were induced by repeated administration of the vitamin D analogue MC903 to the murine skin prior to fungal association [39]. WT *M. sympodialis* robustly colonized the murine skin by day 4 post-infection, with higher loads in the atopic dermatitis-like skin than in control skin, as previously shown [38] (**Figure 5A**). Importantly, under high pH, but not low pH conditions, colonization levels of both, the *rim101*Δ [KPY34] and *rra1*Δ [KPY36] mutants were reduced compared to those of the WT control strain, although differences did not reach statistical significance (**Figure 5A**). The effect was more pronounced at day 7, when the lack of *RIM101* and *RRA1* clearly impaired fitness and resistance to clearance of *M. sympodialis* in skin with elevated pH (**Figure 5B**). Overall reduced fungal burden on day 7 vs. day 4 is consistent with previous observation that *Malassezia* skin colonization is transient in experimental mice [24, 38]. The diminished capacity of the *rim101*Δ and *rra1*Δ mutants to colonize the atopic dermatitis-like murine skin was not due to an intrinsic growth defect of these strains or a defect to initially establish cutaneous colonization, as they colonized the skin equally well at low pH (**Figure 5A**). The difference in skin colonization levels between WT, *rim101* and *rra1* mutant strains did not impact the degree of inflammation of the atopic dermatitis-like skin, as assessed by quantification of ear swelling (**Figures S2A, B**). Together these data demonstrate the importance of *Ms* Rim signaling in fungal survival at alkaline pH, both in vitro as well as in physiologically relevant sites for this commensal microorganism.

**Figure 5.**
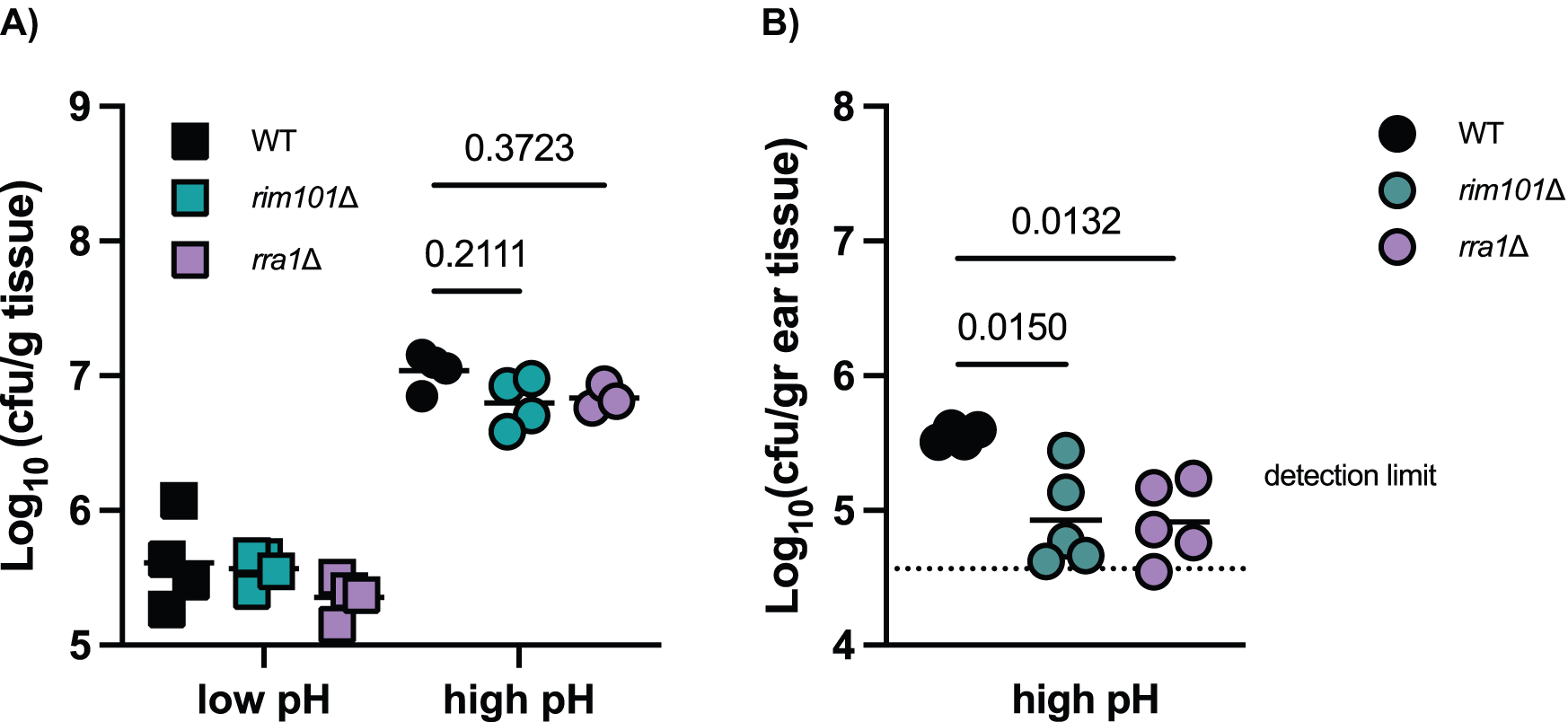
The Rim/Pal1 pathway impacts *M. sympodialis* fitness in a murine model of atopic dermatitis. The ear skin of WT C57BL/6 mice was repeatedly treated with either an ethanol solvent control (low pH, panel A) or with MC903 (high pH, panels A and B) for 10 days to induce an atopic dermatitis-like state. Treated skin was then associated dorsally with *M. sympodialis* WT, *rim101*Δ [KPY34] and *rra1*Δ [KPY36]. The skin fungal load was quantified after 4 days (**A**) or 7 days (**B**) of colonization. Each symbol represents one animal. The mean of each group is indicated (dotted line, detection limit). Two-way ANOVA (**A**) or One-way ANOVA (**B**) were used to determine the statistical significance of the mean colony-forming units (cfu) of each mutant against WT *M. sympodialis*. *, p < 0.05.

## Discussion

*Malassezia* species are among the most common commensal fungi present on the skin, and they are increasingly noted as frequent components of the human gut microbiome [51]. Although most skin sites have a relatively acidic pH, the pH of skin can vary dramatically based on health and disease states. Similar to other fungi, *Malassezia* species possess genes predicted to encode the major components of the pH-responsive Rim signal transduction pathway. We have shown that an intact Rim pathway is required for survival on neutral to alkaline pH as well as elevated salt concentrations, similar to other fungal species. We have also demonstrated that the *M. sympodialis* Rra1 protein is required for fungal survival at alkaline pH, suggesting that Rra1 orthologs are conserved pH-responsive upstream components of Rim signaling in basidiomycetes, likely serving a similar function as the ascomycete Rim21 pH sensors [18]. Finally, we provide evidence that the Rim101/Rra1 pathway increases the fitness of *Malassezia in vivo* in skin exhibiting an elevated pH as it is the case in atopic dermatitis.

Our RNA-Seq analysis of the *M. sympodialis rim101*Δ and *rra1*Δ mutants demonstrated strikingly similar patterns of transcription, further functionally linking *Ms*Rim101 and *Ms*Rra1. Genes demonstrating similar patterns of transcriptional regulation by both Rim101 and Rra1 include those encoding membrane transporters and proteins involved in cell surface adaptation and intracellular trafficking. Proteins involved in similar cell processes are regulated by the Rim101/PacC protein in other fungal species, including the related basidiomycete *C. neoformans* [52]. The fungal cell wall undergoes dramatic changes in structure in response to increases in pH [17], resulting in enhanced fungal survival in this new environmental condition. Therefore, defects in Rim signaling, and the associated failure of pH-responsive cell wall adaptations, may be a major reason for the alkaline pH growth sensitivity in Rim pathway mutant strains.

We also observed Rim pathway-dependent transcriptional changes in genes involved in membrane lipid biosynthesis. This observation is consistent with recent studies in *C. neoformans* Rim signaling in which the phospholipid asymmetry and composition of membranes affect Rim pathway activation [52]. More detailed analysis of the *M. sympodialis* transcriptome is limited by the incomplete annotation of the recently assembled *M. sympodialis* genome [25], with many of the genes demonstrating Rim pathway-dependent expression being listed as uncharacterized proteins. However, these data further demonstrate how the fungal-specific Rim/Pal signaling cascade directs adaptive cellular changes to address the unique challenges resulting from alkaline extracellular environments.

They also suggest that basidiomycetes and ascomycetes have incorporated structurally related but distinct proteins as pH sensors at the plasma membrane.

Interestingly, Rim101 regulation only accounts for a subset of the genes that are up- or down-regulated in response to alkaline pH. However, a large portion of the genes whose expression appears to be Rim101-dependent overlap with the up- or down-regulated genes in the WT in response to DMEM. Overall, these data suggest that many Rim101-regulated processes are conserved between *M. sympodialis* and other distantly related fungal species.

In many fungal species, mutations in Rim/Pal pathway signaling elements result in a marked attenuation of virulence. For example, *C. albicans* Rim pathway mutants are defective in the yeast-hyphal transition in response to elevation in pH, and they are accordingly avirulent in animal models of infection [53]. Related mutations in the *Aspergillus fumigatus* Pal/PacC pathway display reduced hyphal growth in infected lungs [42]. In the case of *C. neoformans*, Rim pathway mutants are defective in many phenotypes typically associated with pathogenesis in this species: these mutant strains are unable to grow well at mammalian pH, as well as in the presence of iron deprivation. Moreover, *C. neoformans rim* mutants fail to incorporate capsular polysaccharide on the cell surface [10]. Consistent with these *in vitro* observations, *C. neoformans rim* mutants have reduced fungal burdens as assessed by quantitative cultures of infected lungs in animal models of cryptococcosis [10]. Paradoxically, mice infected with these attenuated *rim* mutant strains display decreased survival compared to mice infected with wildtype strains [10, 18]. This observation has been explained by excessive immunopathology in the *rim* mutant infections. *C. neoformans* Rim pathway mutations result in an unmasking of typically hidden and immunogenic cell wall epitopes, leading to hyperstimulation of innate immune cell activation and accelerated tissue damage [18].

We did not observe Rim pathway-dependent changes in the degree to which *M. sympodialis* activates macrophages *in vitro*. The *Ms* wildtype, *rim101*Δ, and *rra1*Δ strains induced similar levels of TNF production during *in vitro* macrophage/fungal co-culture experiments. However, in contrast to *C. neoformans* in which the wildtype strains effectively suppress macrophage activation, *M. sympodialis* strains display constitutively high levels of macrophage TNF production, similar to *C. albicans*. Therefore, *M. sympodialis*, though a common commensal, does not appear to shield itself from immune recognition as efficiently as encapsulated fungi such as *C. neoformans*. To fully understand the role of the Rim/Pal1 pathway in the antifungal response to *Malassezia*, future studies with skin-resident cells and model systems harboring features characteristic of the cutaneous niche will be needed.

Emerging data from microbiome studies suggest that *Malassezia* species are not only common skin colonizers but are also found in the human gut, where the pH varies widely from very acidic in the stomach to more alkaline in the small and large intestines. The presence of *Malassezia* species in these micro-niches has been associated with longer term sequelae of continuous antigenic stimulation, including inflammatory bowel disease [54]. Further exploration of the interaction of this commensal fungus with the immune system may elucidate ways in which *Malassezia* species might contribute to health or disease at specific anatomic sites.

## Acknowledgements

This work was supported by NIH grants AI074677 and AI175711 (JAA), F31A140427 (HEB), 1R21AI168672-01A1 (JH and SLL), a National Defense Science and Engineering Grant to KMP awarded by the US Department of Defense (Office of Scientific Research, 32 CFR 168a) and a Swiss National Science Foundation grant to SLL (310030_189255). RNA-Seq studies were supported by a Pilot Grant from the Duke University School of Medicine to support Shared Resource Facilities (Duke Sequencing and Genomic technologies Shared Resource). KMP was supported by a Duke University Bass Instructional Fellowship. Scanning electron microscopy was performed in part at the Chapel Hill Analytical and Nanofabrication Laboratory, CHANL, a member of the North Carolina Research Triangle Nanotechnology Network, RTNN, which is supported by the National Science Foundation, Grant ECCS-1542015, as part of the National Nanotechnology Coordinated Infrastructure, NNCI. We would like to thank Fiorella Ruchti for establishing the experimental conditions for the *in vivo* experiments.

## Author Contributions

KMP, SLL, JH, GI, and JAA were involved with the conception and design of experiments. KMP, CLT, HEB, JTB, EGI, and GI were involved in direct experimentation and acquisition of the data. All authors participated in the analysis and interpretation of the data, as well as in the writing process.

## Supplemental Figure Legends

**Figure S1. Correlation of gene expression in the *rim101*Δ and *rra1*Δ strains.** The *Ms* WT, *rim101*Δ, and *rra1*Δ strains were incubated for 90 minutes in mDixon medium pH 4, mDixon medium pH 7.5, or DMEM tissue culture medium (pH 7.4). Deep RNA sequencing was performed for each strain at each condition, and the log_2_ fold-change for each gene was calculated for the *rim101*Δ and *rra1*Δ strains compared to the WT strain. Gene expression is highly correlated between *rim101*Δ and *rra1*Δ mutants at pH 7.5 and in DMEM, but not at pH 4. Red points indicate genes with expression in *rim101*Δ and *rra1*Δ that is significantly different from WT (adjusted p-value <= 0.05).

**Figure S2. *M. sympodialis* TNF induction is not dependent on fungal cell viability.** WT, *rim101*Δ (KPY34), and *rra1*Δ (KPY36) *M. sympodialis* mutant strains were incubated for 3 days in DMEM + 10% FBS and PenStrep at 30°C preceding a final incubation at 37°C for 16 hours prior to co-culture. WT (H99) *C. neoformans* cells were incubated for 2 days in DMEM + 10% FBS and PenStrep at 30°C preceding a final incubation at 37°C for 16 hours prior to co-culture. WT *C. albicans* (SC5314) cells were incubated for 1 day in DMEM without serum at 30°C prior to co-culture. Bone marrow-derived macrophages (BMMs) were co-incubated for 3 hours with heat-killed (HK) fungal cells (HK, 1 hour at 65°C). BMMs were co-incubated with fungal cells for 6 hours following a 3-hour LPS priming. All strains were at a multiplicity of infection (MOI) of 10:1, fungal cells : BMMs. TNF levels (pg/ml) were assayed from the co-culture supernatant by ELISA. Data represent means from 6 replicates per strain per condition. One-way ANOVA and Tukey’s multiple comparison test were used to compare means. ****, p < 0.0001; *, p < 0.006. Statistical comparisons were made against WT *M. sympodialis*.

**Figure S3. The *M. sympodialis* Rim101/Rra1 pathway does not impact inflammation in a murine model of atopic dermatitis**. To simulate high and low pH under *in vivo* conditions, the ear skin of WT C57BL/6 mice was treated with MC903 (high pH) or EtOH (solvent control, low pH) and then associated with *M. sympodialis* WT, *rim101*Δ [KPY34] and *rra1*Δ [KPY36]. The treatment schedule is indicated at the top of the graph with solid arrows indicating dorsal and ventral application of MC903 or EtOH, and dashed arrows indicate ventral application only of MC903 or EtOH. The ear thickness of mice was determined at the indicated time points. Each datapoint is the mean +/- SD of 4 (**A**) or 5 mice (**B**), respectively.

